# Ancient metapopulations and extreme sex-biased demography revealed by ABC inference and X-Chromosome diversity in Indo-Pacific reef sharks

**DOI:** 10.64898/2026.03.13.711537

**Authors:** Carolin Dahms, Paolo Momigliano

## Abstract

Reconstructing demographic histories in marine species is challenged by extensive geographic distributions, dispersal barriers, high site fidelity, and sex-biased movement. The grey reef shark *Carcharhinus amblyrhynchos* exemplifies these difficulties: despite limited long-distance dispersal, its Indo-Pacific range spans major oceanographic barriers, leading to competing hypotheses of continuous isolation-by-distance versus hierarchical metapopulation structure. We test these alternatives with a new Approximate Bayesian Computation framework and integrate sex chromosome diversity and differentiation analyses. We find strong support for two ancient, deeply separated Indo-Pacific metapopulations that only recently experienced limited secondary contact. X-chromosome diversity shows extreme reductions relative to autosomes (*Q*_*π*_≈ 0.18–0.30 in the central/western Indian Ocean and 0.37–0.46 elsewhere), and *Q*_*FST*_ estimates far below neutral expectations of 0.75. This signature is markedly stronger than in some other carcharhinids and consistent with a reduced effective size of the X chromosome under strong sex-biased dispersal, likely amplified during sequential colonisation events during the last range expansion. Combined evidence of ancient divergence, reduced genetic diversity and extreme depletion of X chromosome diversity supports designating the Western and Central Indian Ocean populations as Evolutionary Significant Units. Our results demonstrate how integrating simulation-based inference with chromosome-specific diversity resolves complex metapopulation histories and reveal how sex-biased dispersal shapes genomic diversity across evolutionary timescales.

## Introduction

Population connectivity - the extent of dispersal among populations - is one of the main factors shaping genetic differentiation across a species range. Quantifying genetic connectivity, defined as the extent to which gene flow influences evolutionary processes, such as inbreeding, genetic drift and the spread of advantageous mutations, is essential to understand a population’s evolutionary potential, and for the delineation of Management Units and Evolutionary Significant Units (ESUs) for conservation (Lowe & Allendorf, 2010). These units guide conservation priorities towards populations of high vulnerability or those possessing unique evolutionary history and adaptations (Dudgeon et al., 2012; Moritz, 1994). For highly mobile marine species such as sharks, assessing genetic connectivity is particularly challenging due to their vast geographic scale of their distribution (Last & Stevens, 2009), variability in life history traits including longevity, maturation and fecundity (Dulvy et al., 2014; Smith et al., 1998), and behaviours such as female philopatry and sex-biased dispersal (Feldheim et al., 2014). Together, these factors generate substantial variation in connectivity patterns among taxa, underscoring the need for species-level genetic structure assessment, as mismatches between genetic connectivity and conservation units may lower management sustainability (Lowe & Allendorf, 2010).

Although range-wide management has been proposed for several shark populations due to their high degree of panmixia (Junge et al., 2019), many species show stronger population structuring due to their unique life history traits and habitat associations (Bernard et al., 2021; Cortés, 2000; Pember et al., 2020). Coral reef-associated *Carcharhinidae*, most of which are listed as Endangered or Vulnerable on the IUCN red list of threatened species, are characterised by complex meta-population structures throughout their wide distribution ranges spanning from the western Indian Ocean to the Pacific (Boissin et al., 2019; Boussarie et al., 2022; Maisano Delser et al., 2019), which are the results of both past historical events such range expansions (e.g. Maisano Delser et al, 2017, Walsh et al, 2022 and Lesturgie et al., 2023) and current environmental dispersal barriers, such as large oceanic distances (Boussarie et al., 2022; Momigliano et al., 2017).

Another factor shaping genetic diversity across species ranges is thought to be sex-biased dispersal, including reduced individual ranges or natal philopatry, due to diverging female and male fitness needs. In sharks, which tend to have disproportionately higher reproductive investment from females, evolutionary theory predicts male biased migration, while females benefit from proximity to nursery habitats (Bonfil et al., 2005). Evidence for male-biased dispersal and female philopatry has been documented by tagging or more frequently on the basis of discordance between biparentally inherited nuclear markers and maternally inherited mtDNA (see Flowers et al., 2016 for a review). Mito-nuclear discordance has been reported in numerous species: the grey reef shark (*C. amblyrhinchos*, Momigliano et al., 2017), shortfin mako (*Isurus oxyrinchus*, Corrigan et al., 2018), Port Jackson shark (Day et al., 2019), bull shark (*Carcharhinus leucas*, Karl et al., 2011) and others (Flowers et al., 2016). However, these genetic studies have likely overestimated the prevalence of sex bias due to unreliable methods such as low sampling or sequencing resolution (Phillips et al., 2021), as well as the inherent constraints of relying on a single, non-recombining locus (mtDNA) to infer female demographic patterns. Indeed, comprehensive reviews including tagging and genetic data identified philopatric and site fidelity behaviours in both sexes in the majority of publications (Chapman et al., 2015; Flowers et al., 2016). To date, only one study by Laso-Jadart et al. (2025) tested female philopatry rigorously with realistic models of sex-biased dispersal, concluding that the mito-nuclear discordance in white sharks is likely driven by evolutionary forces other than demography, casting further doubt on decades of population genetic inference in elasmobranchs.

In this manuscript, we use a novel multi-population Approximate Bayesian Computation (ABC) framework and sex chromosome information from a newly published shark genome (Dahms et al., 2025) to test hypotheses about historical and contemporary genetic connectivity and the effects of sex-biased dispersal on genetic diversity in the grey reef shark (*Carcharhinus amblyrhynchos*), an endangered coral reef– associated carcharhinid. The species is broadly distributed from the Eastern Pacific to the Western Indian Ocean and Red Sea and has high site fidelity to coral reefs (Espinoza et al., 2015). Previous reduced-representation studies revealed strong genetic structure (F_ST_ ≈ 0.5; Lesturgie et al., 2023; Momigliano et al., 2017) between the Central and Western Indian Ocean and the eastern Indian Ocean–Pacific, while differentiation within the Pacific is much lower (F_ST_ < 0.05). There is debate over whether this range-scale structure reflects a continuous metapopulation across vast distances (Lesturgie et al., 2023) or historical effects of heterogeneous habitat (Boussarie et al., 2022). What is clear is that present-day populations result from range expansions (Lesturgie et al., 2023; Walsh et al., 2022) following which genetic diversity has not yet reached migration-drift equilibrium.

Our first goal was to test whether the Central - Western Indian Ocean and the Eastern Indian Ocean–Pacific form a single continuous metapopulation or two distinct metapopulations with separate evolutionary histories. To address this, we developed a novel ABC framework to compare models of continuous migration and hierarchical population structure with and without current and historical migration across metapopulations. This easily scalable approach extends standard divergence models, such as isolation with migration, secondary contact and strict isolation, into a realistic framework for spatially structured meta-populations.

Secondly, we aimed at determining the effects of sex-bias dispersal on the genetic diversity and genetic differentiation across the species range. While previous studies demonstrated strong mito-nuclear discordance in *F*_ST_ estimates for grey reef sharks, even within region (Momigliano et al., 2017), recent work from Laso-Jadart et al. (2025) cast doubt on the validity of such approach. Another way to test the effects of sex bias on genetic diversity is to focus on sex chromosomes. While Y chromosomes are very small and poorly characterized in sharks (but see Dahms et al. 2025 for an exception), X chromosomes are more consistently assembled. In grey reef sharks, female natal philopatry would increase variance in female reproductive success and reduce female-mediated gene flow, leading to a departure from the neutral Wright–Fisher expectation of a 3:4 ratio between effective population sizes of X chromosomes and autosomes (Charlesworth, 2009):

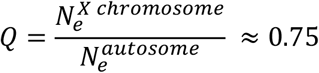

*Q* is therefore a proxy of the Effective Sex Ratio, which can be estimated from the ratio of population genetics statistics - such as nucleotide diversity π and inversely in the fixation index *F*_ST_ - from autosomes and the X chromosome according to the formulas:

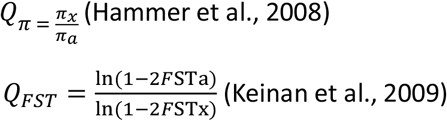

Estimators of the Effective Sex Ratio (ESR) based on nucleotide diversity (*Q*_π_) and population structure (*Q*_FST_) are influenced by sex biases operating over different temporal scales (Emery et al., 2010). Q _π_ is affected by sex bias both before and after population divergence but is primarily shaped by the magnitude of sex bias in ancestral populations. In contrast, *Q*_FST_ is mainly influenced by sex-biased processes occurring after population splits. Comparing these two proxies of ESR can therefore not only reveal whether demographic processes are sex-biased but also highlight contrasting patterns across a species’ evolutionary history (Emery et al, 2010). Although these approaches have previously been used to investigate sex-biased dispersal during human range expansion, they have rarely been applied to non-model organisms. In this study, we tested for the effects of sex-biased dispersal by using a newly published reference genome alongside previously published reduced-representation and whole-genome data from grey reef sharks and other requiem shark species. We estimated *Q* using both methods, evaluated sex-biased demography across temporal scales, examined its relationship with past range expansions, and compared the findings with other species within the same family.

## Methods

### Samples and genotyping

*Carcharhinus amblyrhynchos* DArTseq sequences were retrieved from previous studies (Boussarie et al., 2022; Momigliano et al., 2017), including populations with at least 8 females. Samples covered one location in the Central Indian Ocean from the Chagos archipelago (Chagos, n = 22). Other samples were from the Australian coast of the east Indian Ocean: Ningaloo reef (Ningaloo, n = 23), Rowley Shoals (Rowley, n = 24), and Scott reef (Scott, n = 24). Samples across the West Pacific Ocean included Misool Island in Indonesia (Indo, n = 23) the Southern Great Barrier Reef (South GBR, n = 21), North GBR (n = 19), Chesterfield (n = 30), as well as New Caledonian locations Entrecasteaux (n = 54), Great Northern Lagoon (GLN, n = 51), Grand Astrolabe (n = 46), Matthew (n = 36), Walpole (n = 27. Whole genome sequences (WGS) of an additional population from the Gaafu Dhaalu atoll in Southern Maldives (n = 21) were partially retrieved (Dahms et al., 2025) and partially published in this study utilising the same sequencing approach as per Dahms et al 2025 (Supplementary Table S1). WGS samples were filtered down to sites based on the DArTseq sample with the highest read depth. Another retrieved RADseq dataset (Lesturgie et al., 2023) was analysed separately due to different sequencing approaches. The RADseq data included two populations from the Western Indian Ocean (Juan, n = 13 and Zelee, n = 6), a Coral Sea population (Bampton, n =10), and Central Pacific populations from the Phoenix islands (Niku, n = 21), the remote Palmyra atoll (Palmyra, n = 38) and French Polynesia (Fakarava, n = 27). All sequences were aligned to the grey reef shark reference genome (DDBJ/ENA/GenBank accession number GCA_965287045) using Bowtie2 (v.2.5.0; Langmead & Salzberg, 2012) with *–very-sensitive-local* settings; from the resulting BAM files reads with a mapping quality below 20, coding sequences and repeat elements were removed with samtools (v.1.16.1; Danecek et al., 2021) and bedtools (v.2.27.1; Quinlan & Hall, 2010)

### Sex ratios

We removed coding and transposable elements from all sequences to exclude effects of natural selection, as well as the previously identified pseudo autosomal region within the first 4.9 Mb on the X chromosome (Dahms et al., 2025). Nucleotide diversity (*π*) for autosomes and the X chromosome was estimated separately for each species using non-parametric block bootstrapping across the genome. For each population, we generated 100 bootstraps of one-dimensional site frequency spectra (SFS) in ANGSD (Korneliussen et al., 2014) from Site Allele Frequencies (SAF) using stringent filters that, however, do not affect allele frequencies (-*uniqueOnly 1 -remove_bads 1 -skipTriallelic 1 -minMapQ 30 -minQ 20*) and retaining only sites sequenced for at least 50% of individuals. Because the autosomal and X-linked bootstrap replicates were generated independently, joint replicate pairs were not available. To obtain confidence intervals for the ratio, we therefore treated the empirical bootstrap distributions of πA and πX as independent marginal estimates of their respective sampling distributions. For each species, we computed all pairwise ratios between X-linked and autosomal bootstrap estimates, using the full Cartesian product of bootstrap samples. The resulting empirical distribution of *Q*_*ij*_ was used to derive the median and 97% confidence intervals for *Q*_*π*_. This approach provides a conservative but statistically valid estimate of uncertainty given the absence of paired bootstrap replicates. The same procedure was applied to three previously published carcharhinid sequences: two species known to have female philopatry and sex-biased dispersal (*Carcharhinus limbatus* and *Galeardo cuvier*), and *Carcharhinus acronotus* with unbiased dispersal (Supplementary Table S1). To test for isolation by distance (IBD) of *Q*_*π*_ from their approximate centre of expansion origin in West Papua, we regressed *Q*_*π*_ estimates with least cost geographic distances to the Indonesian location (sampled very close to the center of origin) calculated in R with marmap (v.1.0.12; Pante & Simon-Bouhet, 2013).

Pairwise allelic differentiation, i.e. weighted Wright’s *F*_ST_ for X and autosomes were calculated from two-dimensional SFS in ANGSD for both RADseq datasets. In population pairs with very high *F*_ST_ estimates (> 0.5), the application of Keinan et al’s *Q*_FST_ formula is not possible. Therefore, we transformed estimates into Reynold’s genetic distance: -ln(1-F_ST_), a statistic more sensitive for divergent populations (Libiger et al., 2009; Reynolds et al., 1983), resulting in 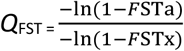. To compare inter- and intraregional *Q*_*FST*_ differences we subdivided the Indian Ocean populations into Western (WIO: Juan, Zelee), Central (CIO: Maldives, Chagos), Eastern (EIO: Rowley, Ningaloo, Scott). The Western Pacific Ocean (WPA) comprised of a Torres Strait population (Indo), Coral Sea populations (South GBR, Bampton), New Caledonia (Entrecasteaux, Grand Astrolabe, Southern NC, Chesterfield, Walpole, GLN); and the remote atoll groups in French Polynesia, Phoenix Islands and Palmyra (Supplementary Table S1). To account for nonindependence of paired *F*_ST_ estimates, we ran 1000 bootstraps with population pair replacement comparing ratios within and between Wregions.

### Runs of homozygosity

ROH were estimated from previously published whole genome resequencing data from 18 individuals: three male and three female grey reef shark sequences per each of the three populations, the Southern Maldives (Gaafu Dhaalu atoll, see previous methods), Ningaloo and Rowley Shoals reefs in Australia (Dahms et al., 2025). The whole genome reads were mapped to the reference assembly using bwa mem (v.0.7.17), samtools and GATK for indel realignment ( v.3.8; McKenna et al., 2010). Variant and invariant sites retrieved with bcftools at *-q 20 -Q 20* were filtered for a minimum read depth of 5 (¼ the average depth), a maximum read depth of 60 (3*average depth). Further filtering in vcftools (v.0.1.17; Danecek et al., 2011) removed indels and retained only bi-allelic sites with 50% missingness. The resulting data consisted of 102,114,956 sites out of the original 1,411,992,160 sites, with a high SNP density of 17kb/SNP. Using plink (v.1.90b6.24; Purcell et al., 2007), we identified runs of homozygosity (ROH) as 100kb regions with the following settings: *--homozyg-snp 100 --homozyg-kb 100 --homozyg-density 50 --homozyg-gap 1000 --homozyg-window-snp 50 --homozyg-window-het 2 --homozyg-window-missing 5 --homozyg-window-threshold 0*.*05*. We kept most settings identical to Stanhope et al. (2023) for comparability, except allowing for smaller regions of >100kb, as well as a relaxed threshold (2) for maximum number of heterozygous sites per window, to control for potential genotype calling errors in our low-coverage data with lower average read depth (~10). For each population we calculated FROH, the proportion of the autosomal genome covered by ROH, and plotted number of ROH (*N*_*ROH*_) against the sum of ROH (*S*_*ROH*_) per individual.

### PSMC

To evaluate demographic history effects on ROH patterns, we estimated past changes in effective population size (*N*_*e*_) for the Southern Maldives, Ningaloo and Rowley Shoals using pairwise sequentially Markovian coalescent (PSMC v.0.6.5; Li & Durbin, 2011). From published sequences with an average coverage of approximately 20X from each of the populations (Dahms et al., 2025), diploid consensus sequences were generated with mpileup according to the PSMC manual and filtered for a minimum read depth of 1/3 the average depth, a maximum read depth of 2 * average depth and a mapping quality > 20. We ran 100 bootstrap replicates with the following parameters: *-N25 -t15 -r5 -p “4 + 25*2 + 4 + 6*”. Time was rescaled with a generation time of 16.4 years and a mutation rate of 8.295 x 10-9 as in Walsh et al. (2022), which is the midpoint of previous estimates for *Carcharhinus galapagensis* from opposite sides of the Central American Isthmus (Maisano Delser et al., 2019). The generation time considers both age of grey reef shark female maturity and the timing of multiple offspring throughout a grey reef shark’s lifetime (Robbins, 2006). The inferred *N*_*e*_ estimates should be considered relative comparisons rather than absolute assessments due to uncertainties of mutation rate estimates.

### ABC Simulations

Using a custom ABC inference pipeline, we tested the hypothesis that grey reef shark populations within the Indo pacific consist of either a single continuous population (Lesturgie et al., 2023) or of independently evolving lineages shaped by oceanographic barriers (Boussarie et al., 2022). We coded three competing demographic models (Figure 1), reflecting one continuous metapopulation (Stepping Stone Model, *SST*) as well as two hierarchical models representing two independent metapopulations (*HIER* and *HIER*_SC_). In the *SST* model, a set of 20 demes of size *N*_e_*D* splits simultaneously at time *T*_1_ from an ancestral population of size *N*_e_*A*, and each neighbouring deme exchanges migrants with a symmetric migration rate *m*. In the model *HIER*, an ancestral population of size *N*_e_*A* splits at time *T*_*1*_ in a left and right lineage of sizes *N*_e_*L* and *N*_e_*R*. At times *T*_2_*L* and *T*_2_*R*, the left and right lineages split in two distinct metapopulations with 10 demes each, of sizes *N*_e_*DL* and *N*_e_*DR*, where neighbouring populations are connected by gene flow according to symmetric migration rates *m*_*n*_*L* (migration between neighbours in the left metapopulation) and *m*_*n*_*R (*migration between neighbours in the right metapopulation). Therefore, both time of a potential expansion, local deme size, and migration rates have different parameters in the two metapopulations. The third model, *HIER*_SC_, is identical to HIER with the exception that it allows migration between the two metapopulations - the rightmost deme of the left lineage and the leftmost deme of the right lineage exchange migrant, assuming an asymmetrical migration rates across the migration bridge.

**Figure 1:**
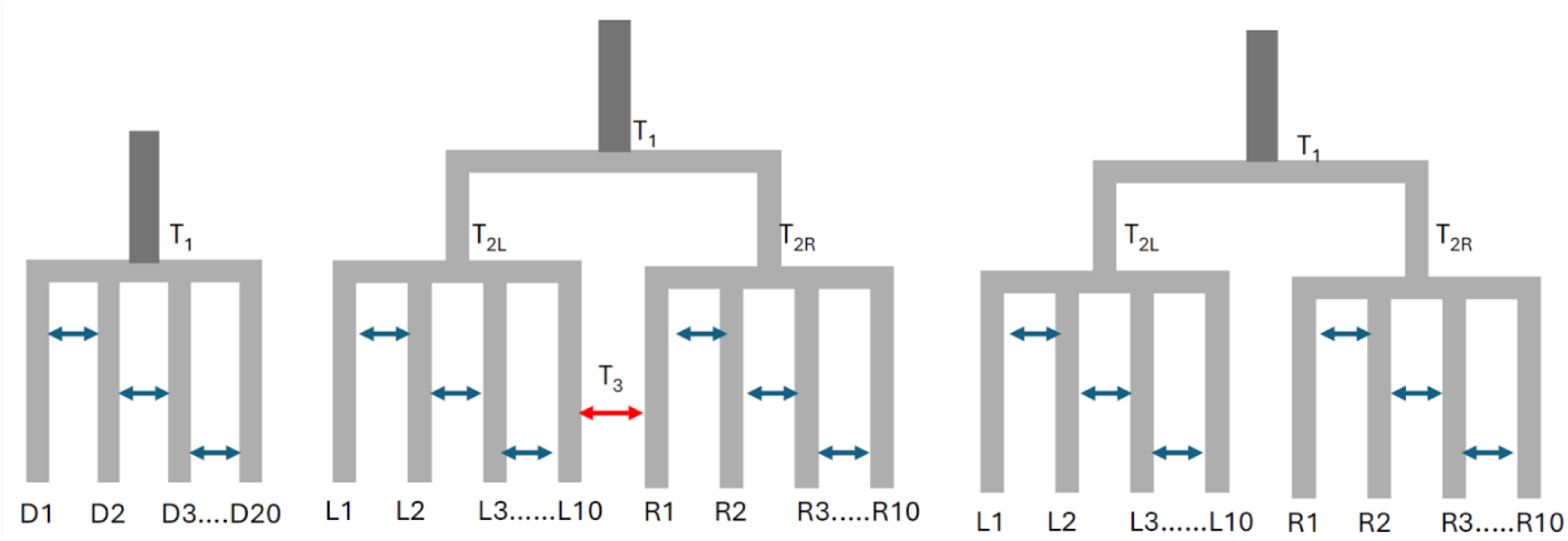
Graphical representation of three demographic models. A simple one-dimensional stepping stone model (SST, left), the simple hierarchical metapopulation model (HIER right), and the hierarchical, secondary contact model (HIER_SC_, center).

We built a custom pipeline (*Build_ABC_refTable*.*py*, https://github.com/Nopaoli/MetapopABC) to simulate these demographic models in *msprime* (Baumdicker et al., 2022), drawing priors for each simulation according to user-specified bounds provided in either a *yaml* or *json* parameter file (Supplementary Table S2). All priors are drawn from a uniform distribution, with the exceptions of *T*_2_*L, T*_2_*R* and *T*_3_. The bounds for these parameters are defined by the user as a fraction of *T*_1_ (in case of *T*_2_L and *T*_2_R) and as a fraction of the lower value of *T*_2_*L* and *T*_2_*R* (in case of *T*_3_), and they are drawn from a beta distribution where the user sets both the alpha and beta parameters. Migration rates are sampled so that they generate a uniform distribution of *N*_e_*m* (though the pipeline also allows to sample directly a uniform distribution of migration rates). After the simulations are run with *msprime*, the pipeline employs *tskit* (Jeffery et al., 2026) to compute summary statistics. We employed a set of summaries from both one and two populations, including Watterson’s Θ, π (average number of pairwise differences), Tajima’s D, and the frequencies of each folded Site Frequency Spectrum (SFS) bin, standardized by the full sequence length. For each pair of sampled populations, we computed Hudson’s *F*_ST_ (Hudson et al., 1992), as well as absolute (*d*_xy_) and net (*d*_a_) divergence (Nei & Li, 1979). All observed summary statistics fell within the distribution of simulated estimates (Supplementary Figures S1-5). The full pipeline and a detailed user guide are available on GitHub (https://github.com/Nopaoli/MetapopABC), and an annotated yaml file containing all parameters used for the simulations is included in the Supplementary Table S2. The pipeline is fully customizable, including the number of demes in each metapopulation.

We simulated 10,000 datasets from each scenario, using a mutation rate *μ*=1.9e^-8^ (as in Lesturgie et al. 2023), and simulating loci that replicate our empirical sequence length and locus structure, i.e. 71 500 of a length of 70 bp, collecting five diploid individuals from each sampled deme. We designed a deme sampling scheme to approximate as closely as possible the actual geographic distances among our sampling locations. For Model SST, we selected demes D2, D14, and D16, corresponding respectively to our Chagos, Ningaloo, and Indonesian samples. The rationale was to include one population closer to the edge of the distribution (Chagos) and two at the opposite end, which are geographically closer to each other. For Models HIER and HIERSC, we instead sampled the central deme of the left metapopulation (Chagos) and two demes from the right metapopulation (R3 and R6). Recognizing that the spacing within Model SST is somewhat arbitrary—and acknowledging that this could influence both model choice and parameter estimation—we conducted additional analyses using only two populations. Specifically, we used D2 and D14 for SST, and L5 and R3 for *HIER* and *HIER*_SC_. This allowed us to test whether arbitrary population spacing was biasing model choice.

Model choice was performed using an ABC random forest approach (ABC-RF; Pudlo et al., 2016), as implemented in the *abrcrf* R package. We processed the reference table of 10,000 datasets per scenario and ran 10 independent replicate forests (distinct random seeds) to assess consistency, each with 1,000 trees. We first preformed model selection using a hierarchical ABC approach (see Momigliano et al. 2017): first testing SST vs. HIER, to determine whether a single metapopulation or a set of hierarchical metapopulations best fit our data. Secondly, we compared the winning model HIER vs. HIERSC, to test whether there was support for ongoing migration between the two metapopulations. We preformed model choice selection for all three models simultaneously. Once the best scenario was identified, we simulated 40,000 more datasets and estimated parameters from a reference table of 50,000 simulations, also using an ABC-RF approach (Raynal et al., 2019). We checked that our prior distributions were generating summary statistics consistent with our data by plotting the distribution of every summary statistic and consulting a Linear discriminant Analyses (LDA, Supplementary Figure S6). We computed NMSE (Normalized Mean Square Error) as well as Coefficient of determination (Q^2^; Raynal et al., 2019) to assess identifiability of each parameter.

## Results

Nucleotide diversity differed across regions and chromosomes, with X chromosomes showing two to five-fold reductions compared to autosomes and being lowest in the West-Central Indian Ocean *C. amblyrhynchos* populations (Figure 2B, Supplementary Table S3). It should be noted that *π* estimates across different reduced representation sequences (RADseq vs DarTseq) can have slight variation due to distinct ascertainment bias, yet ratios between X and autosomes should not be strongly affected. Accordingly, *π* X:A ratios were extremely below the neutral expectation (*Q*_*π*_ < 0.75) across the *C. amblyrhynchos* distribution range (Figure 2), in line with other carcharhinids known to exhibit sex-biased dispersal (*C. limbatus* and *G. cuvier*), but below *C. acronotus* at demographic neutrality (Figure 2B). The lowest ratios were in the Western and Central Indian Ocean populations, with medians ranging from 0.18 to 0.30, which were negatively correlated with least cost distance to Indo, the approximate location of past range expansion (Pearson’s *r* = −0.81, t(5) = −3.07, p = 0.03; Supplementary Figure S7). *Q*_*π*_ estimates were higher in the Eastern Indian Ocean and the Western Pacific (0.37–0.46), the latter of which followed no significant spatial correlation (Pearson’s *r* = −0.34, t(10) = −1.15, p = 0.27).

**Figure 2.**
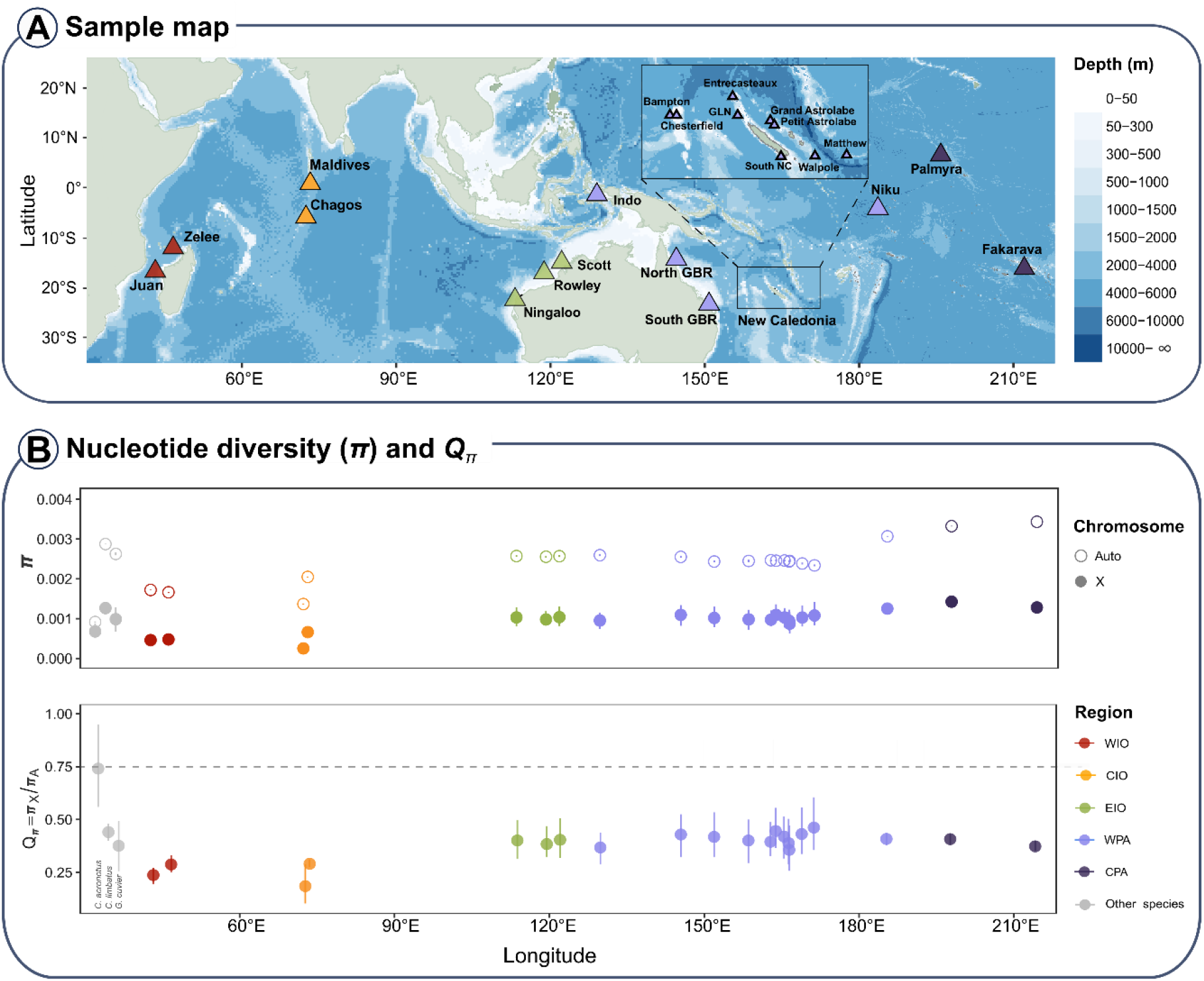
Sampling locations (A) and Q_π_ ratios (B). A: Locations from both RADseq and DArTseq datasets spanning the Western Indian Ocean (WIO, red), Central Indian Ocean (CIO, orange), Eastern Indian Ocean (EIO, green), Western (WPA, light purple) and Central Pacific Ocean (CPA, dark purple). Locations of reference species (Carcharhinus acronotus, Carcharhinus limbatus, Galeocerdo cuvier) in grey are not shown. B: bootstrapped nucleotide diversities are shown as medians and 97% Confidence intervals for autosomes and X chromosomes, with the bottom panel showing the X:autosome ratios across longitudes. The dashed line at 0.75 represents the neutral expectation, with values below indicating X chromosome Ne reduction consistent with male biased dispersal.

Allelic differentiation *F*_ST_ was highest when comparing Western and Central Indian Ocean populations to other regions, with autosome *F*_ST_ ranging from 0.31 to 0.44, while X chromosomes showed even higher differentiation, ranging 0.47 to 0.66 (Figure 3). Among the Eastern Indian Ocean and Western Pacific, *F*_ST_ A ranged from around 0.006 to 0.048, while X*F*_ST_ included values up 0.09 (Figure 3). The median A:X *F*_ST_ ratio (*Q*_*FST*_) was below the neutral expectation of 0.75 for both WIO and CIO datasets (0.63 and 0.54, respectively), estimates ranging as low as 0.13, with some outliers > 1 (Figure 3, Supplementary Figure S8). We found a strong regional effect on *Q*_*FST*_ –the WIO dataset indicated a noticeable higher *Q*_*FST*_ slightly above demographic neutrality (median = 0.92; 97% CI [0.83; 1.05]) in population pairs stemming from the same oceanic regions than those from different regions (median = 0.59, CI [0.53;0.63]); Supplementary Table S4).

**Figure 3:**
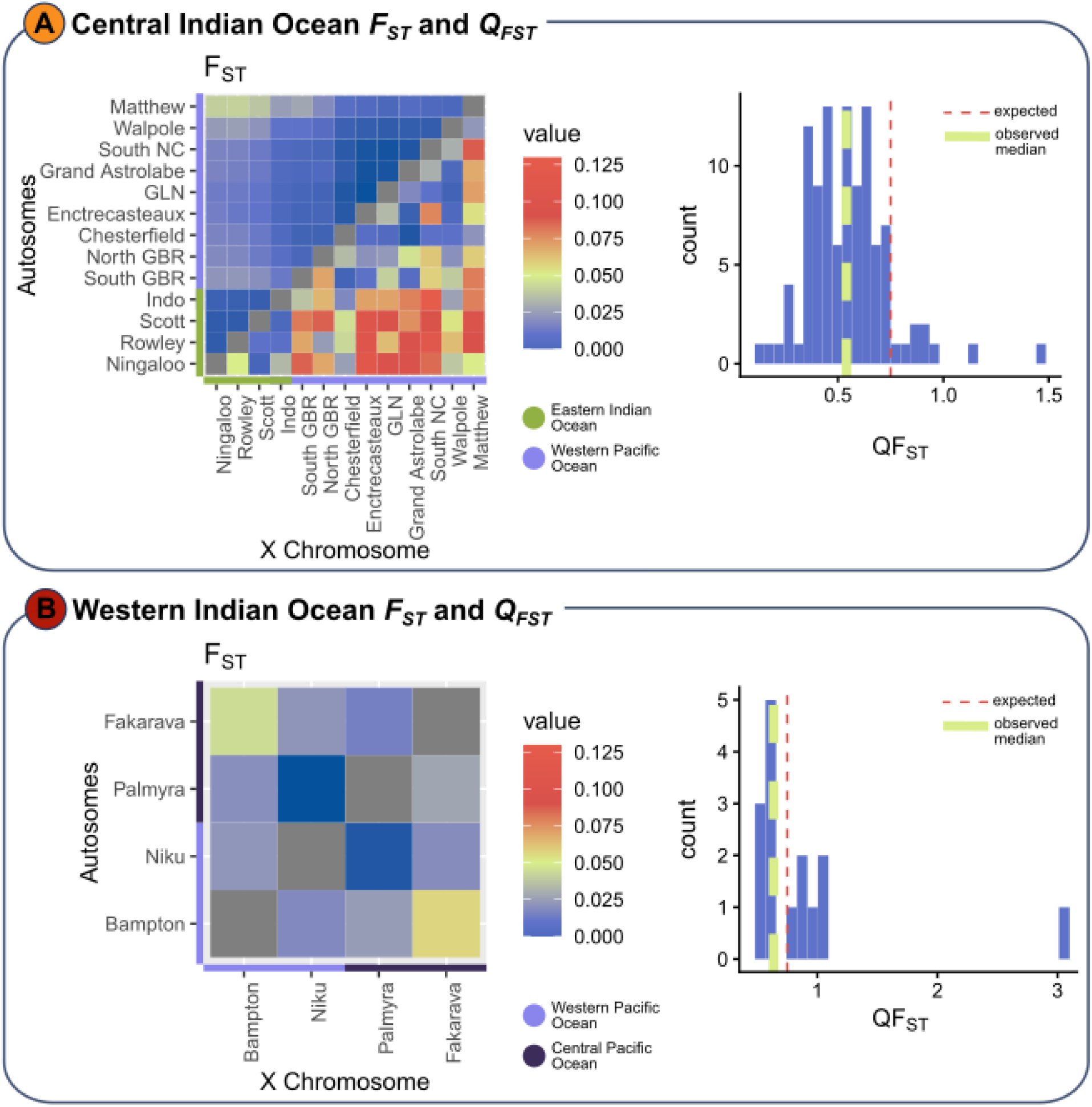
F_ST_ matrices and histograms of Q_FST_. Autosomal and X chromsome Fsts (left) and histogram of Q_FST_ frequencies (right) are shwon for both datasets – DArTseq (a) and Radseq(b). Indian Ocean populations (Fst > 0.3) are removed from Fst matrices for clarity. In histograms, the expected Q_FST_ at netural demography of 0.75 is indicated by the dotted red line line, while the green line shows the observed median.

*N*_*e*_ trajectories showed clear divergence since around 1 million years ago, as *N*_*e*_ increased in Rowley Shoals and Ningaloo populations, reaching *N*_*e*_ estimates of around 13-15 thousand, while a sustained decline in the Maldives resulted in an around five-fold lower *N*_*e*_ at present (Figure 4A). Tests of runs of homozygosity (ROH) patterns were consistent with histories of effective population sizes (Figure 4B)., and indicated intermediate levels of inbreeding compared to other species (Supplementary Figure S9). The median ROH length was relatively short at 124kb, the longest reaching 746kb. We identified 3,254, 1,496 and 1,059 ROH(>100kb) for the Maldivian population (n = 6), and two Western Australian populations, Ningaloo (n = 6) and Rowley Shoals (n = 6), respectively, no ROH longer than 1 Mb were detected (Figure 4B). FROH(>100kb), i.e. the proportion of the genome made up of identified ROH, was 15.2%, 7.4% and 5% for the Maldives, Ningaloo and Rowley Shoals, respectively.

**Figure 4:**
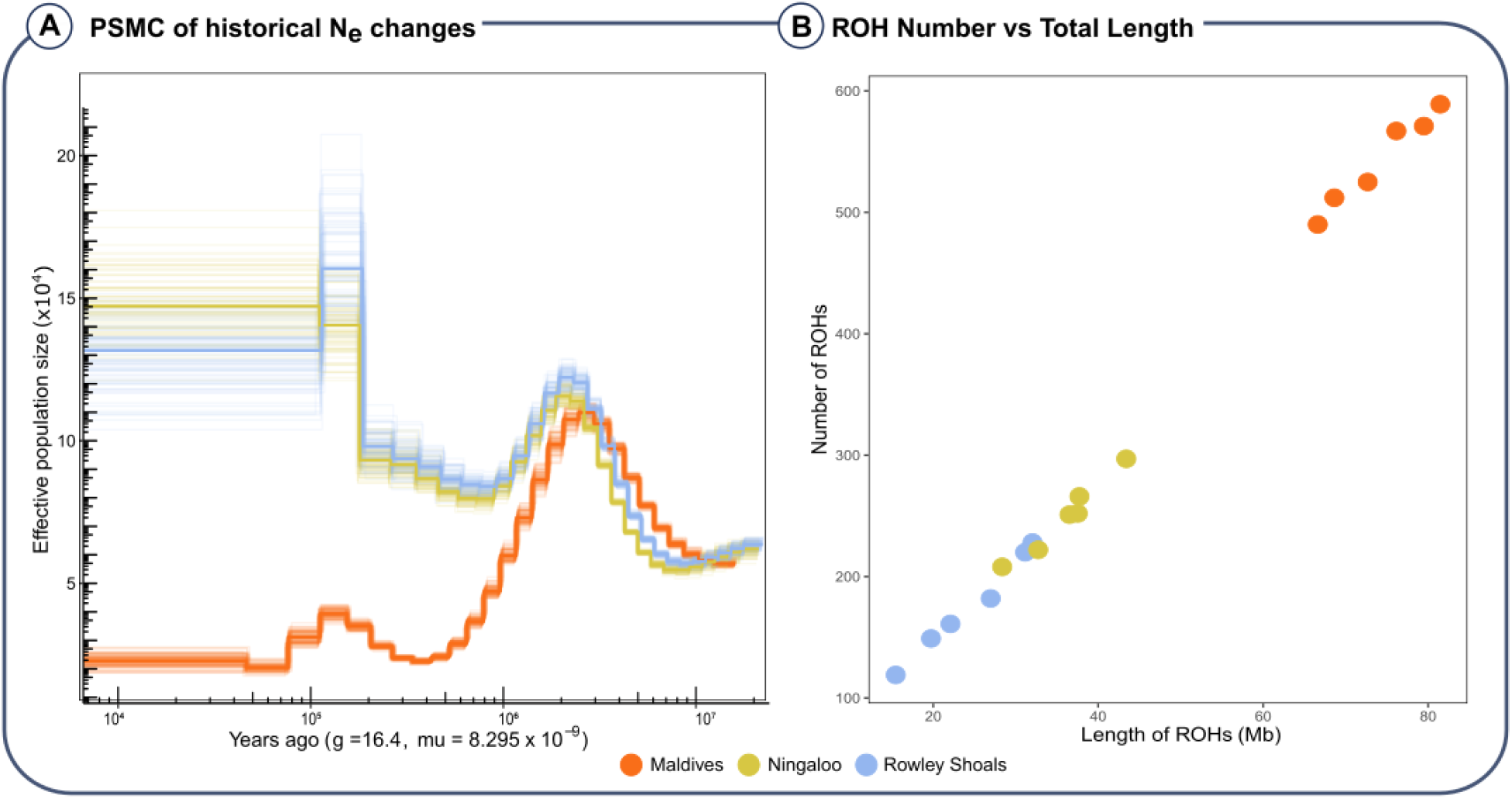
**(A) Historical effective population sizes (Ne) inferred with PSMC**. Ne changes calculated from individual whole genome sequences from three populations – the Maldives (orange), Ningaloo (yellow), and the Rowley Shoals (blue) - were scaled with a generation time (g) of 16.4 years, and a mutation rate (mu) of 8.295 x 10^-9^. The bright lines represent the median from 100 bootstraps (lighter lines). **(B) Relationship between number and sum of ROH lengths (Mb)** per individual from Maldives (orange, n = 6), Ningaloo (yellow, n = 6) and Rowley Shoals (blue, n = 6), including ROH > 100kb across all chromosomes.

ABCF-RF model choice shows unambiguous support for a hierarchical metapopulation structure, whereby Chagos belongs to a distinct metapopulation than Ningaloo and Indonesia (Table 1). The posterior probability for the *HIER* model when compared to the SST model was 0.985 when three demes were sampled, and 0.984 when only one deme per metapopulation was sampled, suggesting strong support for a hierarchical metapopulation structure, and little influence of arbitrary spacing between the sampled demes on model choice. Similarly, the posterior probability of *HIER*_SC_ was 0.971 (when compared with *HIER*) and 0.976 (in the three-model comparison) when three demes were sampled, and 0.962 and 0.986 when two demes were sampled, indicating very strong support for secondary contact among the two metapopulations.

**Table 1:** Results from ABC-RF model comparison. Results are averages from 10 independent replicate forests, each with 1000 trees. Models = models compared, Pops = number of sampled demes (either 2 or 3), Best = identified best model, Posterior= posterior probability of the best model, Votes (SST, HIER and HIER_SC_) represent the number of votes for each model, Prior Error= prior error rate estimated from out-of-bag (OOB) predictions.

| Models | Pops | Nsim | Best | Posterior | Votes SST | Votes HIER | Votes HIER<sub>SC</sub> | Prior Error |
| --- | --- | --- | --- | --- | --- | --- | --- | --- |
| *SST, HIER* | 3 | 20000 | *HIER* | 0.985 | 28 | 972 | NA | 0.0017 |
| *HIER, HIER<sub>SC</sub>* | 3 | 20000 | *HIER<sub>SC</sub>* | 0.971 | NA | 230.9 | 769.1 | 0.09 |
| *SST, HIER, HIER<sub>SC</sub>* | 3 | 30000 | *HIER<sub>SC</sub>* | 0.976 | 2 | 333.2 | 665 | 0.074 |
| *SST, HIER* | 2 | 20000 | *HIER* | 0.984 | 97.8 | 902.2 | NA | 0.009 |
| *HIER, HIER<sub>SC</sub>* | 2 | 20000 | *HIER<sub>SC</sub>* | 0.962 | NA | 227.4 | 772.6 | 0.09 |
| *SST, HIER, HIER<sub>SC</sub>* | 2 | 30000 | *HIER<sub>SC</sub>* | 0.986 | 11.3 | 275.3 | 713.4 | 0.083 |

Not all parameters were well identifiable (Supplementary Table S5). Generally, the forests had good predictive power for split times (Q^2^ between 0.63 and 0.893), and contemporary deme size (> 0.9), moderate predictive power for N_e_A (0.624) and very low predictive power for the ancestral size of the two metapopulations (*N*_e_*L and N*_e_*R*). Within meta-ppulation migration rates (*m*_*n*_*L* and *m*_*n*_*R*) were clearly identifiable (Q^2^ of 0.77 and 0.74, respectively), while migration rates across the metapopulation bridge were more poorly predicted. The results generally point to strong migration within each metapopulation, and lower and asymmetric migration across the metapopulations following recent secondary contact.

## Discussion

In this study, we combined a custom Approximate Bayesian Computation (ABC) simulation approach with analyses of X-chromosome diversity in non-genic regions to test demographic history scenarios of the grey reef shark across its distribution range and identify sex-biased demographic processes. Using the new grey reef shark reference genome alongside previously published RAD-seq and whole-genome data, we found strong support for hierarchical metapopulation structure, with deep divergence between the Central Indian Ocean on one side and the Eastern Indian Ocean and Western Pacific Ocean on the other, including evidence of recent secondary contact. These results confirm the existence of two distinct, long-isolated lineages across the species’ distribution range and support the hypothesis that deep ocean expanses have acted as a strong dispersal barrier. Estimates of *Q* derived from genetic diversity (*Q*_*π*_) and fixation indices (*Q*_*FST*_) were substantially lower than neutral expectations, providing a consistent signal of male-biased demography across both metapopulations at different time scales. This bias was particularly pronounced for genetic diversity in the WIO and CIO, where a significant *Q*_*π*_ gradient with distance from the origin of expansion is consistent with male-biased dispersal during sequential colonization events. Ultimately, the evidence of deep divergence, reduced genetic diversity, and extreme sex-bias in Western and Central Indian Ocean populations supports their designation as Evolutionarily Significant Units.

The ABCRF confirmed the central Indian Ocean to be part of an ancient metapopulation which split before the Pacific Ocean populations that have experienced low degrees of secondary gene flow. The deep divergence, low genetic diversity is in line with previous analyses showing strong genetic structuring using both nuclear (Boussarie et al., 2022; Lesturgie et al., 2023) and mtDNA markers (Momigliano et al., 2017). The results indicate deep divergence between the two metapopulations, with estimated split times of nearly 80,000 generations (median: 77,898). Assuming a generation time of 16.4 years, this corresponds to approximately 1.28 million years ago, broadly consistent with the divergence in demographic histories inferred from PSMC analyses. However, it is important to note that in structured populations the instantaneous inverse coalescent rate inferred by PSMC does not necessarily reflect changes in census or effective population size, but may also capture shifts in connectivity among subpopulations or simply arise from spatial population structure (Chikhi et al., 2010; Mazet et al. 2016). Consequently, some of the inferred historical *Ne* trajectories may partly reflect the effects of metapopulation structure—and temporal changes in connectivity—rather than population size per se. Nevertheless, these findings suggest that the two metapopulations likely originated from distinct range expansions of ancestral lineages, after which gene flow was recently re-established.

Though estimates of parameters such as deme size and migration rates are inevitably influenced by model specification (e.g., the number of simulated demes and the assumption of a one-dimensional population structure), qualitative comparisons between the two metapopulations reveal clear differences. In particular, demes in the Central Indian Ocean metapopulation are much smaller (median: 1,629) than those in the Eastern Indian Ocean and Pacific metapopulation (median: 5,047). Because absolute migration rates are very similar between the two regions, this difference implies that far fewer migrants (Nm) are exchanged among demes each generation in the Central Indian Ocean metapopulation. Overall, these results suggest that subpopulations in the Central Indian Ocean tend to be smaller and more isolated than those elsewhere in the species’ range. This interpretation is consistent with the substantially lower levels of genetic diversity observed in this region.

Divergent histories are mirrored by distinct estimates of *Q*_*π*_ across the species range. While *Q*_*π*_ estimates are always far below the expectation of 0.75 across populations (0.15-0.46), the strongest departures are recorded in the Chagos and Juan populations (0.18, 0.24, respectively) suggesting extremely reduced X chromosome Ne in the Central and Western Indian Ocean. The highest values recorded from *C. amblyrhynchos* were comparable to other carcharhinids known to exhibit female philopatry and sex-bias dispersal, such as *G. cuvier* (0.37) and *C. limbatus* (0.44), and much lower than *C. acronotus*, which has no clear documentation of sex bias dispersal and exhibits a ratio within the neutral expectation of 0.75 (Dimens et al., 2019). The more extreme *Q*_*π*_ could also be caused by a recent reduction in population size, such that the population size for the X chromosome has approached its new equilibrium faster than that for the autosomes (Pool & Nielsen, 2007) which is consistent with Ne reductions in the Indian Ocean. Interestingly, Qpi in the Indian Ocean decreases with distance from the approximate centre of the last expansion origin (in West Papua, Parson’s *r* = −.81, p = 0.03), which could indicate male biased colonisation particularly in the remote and deeply diverged Indian Ocean populations. This suggests significantly higher drift experienced by the X chromosome at some point or throughout the time of expansion waves, leading to reduced X Ne and high *Q*_*π*_. This has been observed in humans during their out of Africa migration, where both X chromosome drift due to male biased colonisation and strong selection on X has been proposed (Keinan et al., 2009).

Similarly, *Q*_*FST*_ were below neutrality expectations across most of the population comparisons (Figure 3). High *F*_ST_ in in both autosomes and X chromosomes were in line with bathymetry being a dispersal barrier for both males and females as proposed by previous studies (Momigliano et al., 2017). When comparing inter and intra-regional effects, *Q*_*FST*_ was lower on smaller spatial scales for the CIO dataset (Figure 3). A lower intra-regional ratio agrees with the hypothesis that within regions, genetic structure is shaped by higher female philopatry, as indicated by manyfold higher mtDNA structuring on smaller regional scales, where male migrations are more likely between reefs along continental shelves. This contrasts with comparisons across larger distances: in comparisons involving remote populations like Chagos, Maldives or Walpole, *Q*_*FST*_ was higher compared to most other comparisons, as both male and female dispersal is limited equally resulting in more homogenous differentiation in X and autosomes. The opposite observation in the WIO dataset, with higher ratios > 0.75 within regions in the Pacific Ocean than between IO and PO is surprising. It could be explained by very low *F*_STa_ and *F*_STx_ estimates in some PO comparisons, involving populations like Fakarava and Palmyra, which increases ratio estimates (Supplementary Figure S8). It may also indicate other factors not accounted for - demography alone has not held up as a reliable predictor of departures from X:A expectations in many species including humans (Hammer et al., 2008; Keinan et al., 2009) and primates (Osada et al., 2021). Often rather than philopatry, evolutionary factors such as natural selection have been identified as significant drivers, such as in Drosophila, where X alleles in hemizygous males are more likely to experience selective pressures (Andolfatto et al., 2011). In primates and humans, negative selection against X chromosome during nuclear swamping in admixing populations (Osada et al., 2013) or repeated selective sweeps on across evolutionary timescales have been proposed (Skov et al., 2023). Although in our analyses strong X:A ratio departures remained after excluding nongenic regions supporting sex biased dispersal, these were limited to the resolution of reduced representation data. Called genotypes did not include any sites on the Y chromosome and limited number of polymorphisms on X, which prohibited our ability to find selection outliers, and test other potential factors like background selection and hitchhiking effects on the sex chromosomes, which may illuminate further patterns (Evans et al., 2014).

Significantly larger numbers of longer ROH were observed in the Maldivian population compared to Western Australian populations in the Eastern Indian Ocean, in line with a lower effective population size in the remote atolls of southern Maldives, which experienced historical effective population declines (Figure 4A). Compared to previously reported ROH for other shark species, the grey reef shark has a higher number of ROH than the endangered whale shark (*Rhincodon typus*), brown banded bamboo shark (*Chiloscyllium punctatum*, IUCN: near threatened) and the cloudy catshark (*Scyliorhinus torazame*, IUCN: least concern). Only the Southern Maldivian grey reef shark population had ROH counts higher than those of the white shark and approaching estimates of the inbred shortfin mako, although shorter in length and with no ROH of >1Mb being detected (Supplementary Figure S9). FROH did not reach levels as those observed in other notoriously inbred wild populations such as the Devils hole pupfish (Tian et al., 2022) or the hammerhead shark (Stanhope et al., 2023), however the Maldivian FROH (>100kb) was akin to estimates of the shortfin mako at 15% FROH (>100kb; Stanhope et al., 2023). The larger number of relatively small size ROH in *C. amblyrhynchos* reflects historical long-term reduced *N*_*e*_ rather than recent inbreeding or severe bottlenecks. This is important for two reasons: first, it suggests that the observed variance in genetic diversity across the grey reef shark distribution range is likely not due to the recent, ever so dramatic population declines, but rather a reflection of a broad-scale metapopulation structure, historical isolation, and perhaps distinct histories of range expansions and contractions, as supported by the PSMC trajectories (Figure 4A) and ABC inference; and second, the lack of inbreeding depression (i.e. long ROH) suggests that fitness should remain relatively high despite the observed low genetic diversity (Kyriazis et al. 2025).

Our findings have significant implications for *C. amblyrhynchos* conservation management. The WIO and CIO populations can be designated as Evolutionary Significant Units due to convergent evidence of deep mito-nuclear differentiation from other regions, independent evolutionary trajectories, and limited gene flow (determined by demographic modelling) as well as reduced genetic diversity with extreme depletion on the X chromosome. Although the lack of long ROH may suggest absence of significant inbreeding depression, their stark isolation may aggravate local population declines that cannot be rescued by immigration in the future if critical habitat continues to be mismanaged. Currently, 7.1% of critical chondrichthyan habitat is within designated MPAs and only 1.2% within fully protected no-take areas in the WIO (Cochran et al., 2026). Besides collaborative large-scale management, regional decision making would benefit from incorporating philopatry effects to account for disproportionate genetic vulnerability of females and inform spatial management to protect vital breeding and feeding grounds.

## Data availability

The pipeline for ABC analyses is available at https://github.com/Nopaoli/MetapopABC.

