## Supplementary Materials for "Ancient metapopulations and extreme sex-biased demography revealed by ABC inference and X-Chromosome diversity in Indo-Pacific reef sharks"

***Supplementary Table S1: Sample information.*** *For each population, their location and oceanic region, number of males and females (N), sequence data type, as well as citations and sequence accession numbers are given.*

| *Population* | *Location* | *Region* | *N (Females \| Males)* | *Data type* | *Reference* | *Accession* |
|---|---|---|---|---|---|---|
| *Chagos* | Chagos | Central Indian Ocean | 10 \| 12 | DArTseq | Dahms et al., 2025 | PRJEB90057 |
| *Maldives* | Southern Maldives | Central Indian Ocean | 12 \| 8 | WGS | Dahms et al., 2025 | PRJEB90057 |
| *Ningaloo* | Ningaloo Reef | Eastern Indian Ocean | 18 \| 5 | DArTseq | Momigliano et al., 2017 | PRJNA795958 |
| *Rowley* | Rowley Shoals | Eastern Indian Ocean | 14 \| 10 | DArTseq | Momigliano et al., 2017 | PRJNA795958 |
| *Scott* | Scott Reef | Eastern Indian Ocean | 13 \| 11 | DArTseq | Momigliano et al., 2017 | PRJNA795958 |
| *Chesterfield* | Coral Sea | Western Pacific Ocean | 24 \| 16 | DArTseq | Boussarie et al., 2022 | PRJNA795958 |
| *Entrecasteaux* | New Caledonia | Western Pacific Ocean | 28 \| 26 | DArTseq | Boussarie et al., 2022 | PRJNA795958 |
| *GLN (Great Northern Lagoon)* | New Caledonia | Western Pacific Ocean | 18 \| 33 | DArTseq | Lesturgie et al., 2023 | PRJNA795958 |
| *Grand Astrolabe* | New Caledonia | Western Pacific Ocean | 31 \| 15 | DArTseq | Boussarie et al., 2022 | PRJNA795958 |
| *Indo* | Torres Strait | Western Pacific Ocean | 20 \| 3 | DArTseq | Momigliano et al., 2017 | PRJNA795958 |
| *Matthew* | New Caledonia | Western Pacific Ocean | 15 \| 21 | DArTseq | Boussarie et al., 2022 | PRJNA795958 |
| *North GBR (Great Barrier Reef)* | Coral Sea | Western Pacific Ocean | 13 \| 6 | DArTseq | Momigliano et al., 2017 | PRJNA795958 |
| *South GBR (Great Barrier Reef)* | Coral Sea | Western Pacific Ocean | 12 \| 9 | DArTseq | Momigliano et al., 2017 | PRJNA795958 |
| *South NC (New Caledonia)* | New Caledonia | Western Pacific Ocean | 12 \|27 | DArTseq | Boussarie et al., 2022 | PRJNA795958 |
| *Walpole* | New Caledonia | Western Pacific Ocean | 18 \| 9 | DArTseq | Boussarie et al., 2022 | PRJNA795958 |
| *Fakarava* | French Polynesia | Central Pacific Ocean | 13 \| 14 | RADseq | Lesturgie et al., 2023 | PRJNA917473 |
| *Niku* | Phoenix Islands | Central Pacific Ocean | 12 \| 9 | RADseq | Lesturgie et al., 2023 | PRJNA917473 |
| *Palmyra* | Central Pacific Ocean | Central Pacific Ocean | 19 \| 19 | RADseq | Lesturgie et al., 2023 | PRJNA917473 |
| *Juan* | Western Indian Ocean | Western Indian Ocean | 8 \| 5 | RADseq | Lesturgie et al., 2023 | PRJNA917473 |

| | | | | | | |
|---|---|---|---|---|---|---|
| *Zelee* | Western Indian Ocean | Western Indian Ocean | 4 \| 2 | RADseq | Lesturgie et al., 2023 | PRJNA917473 |
| *Bampton* | Coral Sea | Western Pacific Ocean | 10 \| 0 | RADseq | Lesturgie et al., 2023 | PRJNA917473 |

***Supplementary Table S2: YAML file containing all simulation parameters for ABC inference.***


```
# ========================
# ABC reference table config
# ========================
# ---- Core simulation design ----
mu: 1.9e-8          # [REQUIRED] Mutation rate per bp per generation (float)
recomb_rate: 0.0      # NEW: Recombination per bp per generation (rho). Set >0 to enable recomb within loci.
reps: 71500          # Number of independent loci per draw (int >=1)
length: 70        # Base pairs per locus (float >0)
seed: null          # Base RNG seed (int) or null for random
from_sfs: false       # If true, compute ALL stats from SFS (no variances)
variance: false       # If true (and from_sfs=false), compute across-locus variances (slower, larger outputs)
no_singleton: false    # If true, drop MAC1 bin from folded 1D SFS outputs
# ---- Parallelism ----
jobs: 45             # Number of worker processes (int >=1)
workers: null        # Placeholder for future msprime num_threads (currently unused)
# ---- Metapopulation sizes (deme counts) ----
m1_n: 20            # Total demes in Model 1 (1D chain)
m2_nl: 10           # Left demes in Models 2 & 3 (1D chain)
m2_nr: 10           # Right demes in Models 2 & 3 (1D chain)
# ---- Sampling (choose up to 5 demes per model) ----
# Provide comma-separated labels:
m1_pops: "P2,P14,P16"   # Sampled demes for Model 1
m2_pops: "L5,R3,R6"     # Sampled demes for Models 2 & 3 (same sampling used for Model 3)
# ---- Time priors ----
prior_t1_min: 1.0e2     # T1 ~ Uniform[min, max]  (generations)
prior_t1_max: 5.0e5
# T2 and T3 are drawn as *fractions*, then mapped:
t2_frac_min: 0.1       # lower bound for T2/T1 mapping
t2_frac_max: 0.9       # upper bound for T2/T1 mapping
t3_frac_min: 0.1       # lower bound for T3/min(T2L, T2R) mapping (Model 2 only)
```

*t3_frac_max: 0.9 # upper bound for T3/min(T2L, T2R) mapping*

*t2_beta_alpha: 2.0 # Beta(alpha, beta) for T2 fractions*

*t2_beta_beta: 1.0*

*t3_beta_alpha: 3.0 # Beta(alpha, beta) for T3 fraction (Model 2 only)*

*t3_beta_beta: 1.2*

*# ---- Migration priors (probabilities in m-space) ----*

*# use_loguniform_m: true*

*# Symmetric nearest-neighbor migration within each 1D chain:*

*# Model 1 (NEW keys for the new script):*

*# m_neighbor1_min: 1.0e-5*

*# m_neighbor1_max: 1.0e-1*

*# Model 2 within chains (Left / Right):*

*# m_neighborL_min: 1.0e-5*

*# m_neighborL_max: 1.0e-1*

*# m_neighborR_min: 1.0e-5*

*# m_neighborR_max: 1.0e-1*

*# ASYMMETRIC bridge (Model 2 only), L(last) -> R0 and R0 -> L(last) at T3:*

*# m_bridge_L2R_min: 1.0e-6*

*# m_bridge_L2R_max: 1.0e-2*

*# m_bridge_R2L_min: 1.0e-6*

*# m_bridge_R2L_max: 1.0e-2*

*# ---- Optionally draw migration from Nem (expected number of migrants per generation) ----*

*# If true, within-chain m for Model 1 and Model 2 are derived from Nem / Ne_deme (symmetric within each chain).*

*draw_within_from_Nem: true*

*use_loguniform_Nem: false*

*# Model 1 within-chain Nem prior (if draw_within_from_Nem: true)*

*Nem_neighbor1_min: 1*

*Nem_neighbor1_max: 100*

*# Model 2 within-chain Nem priors (if draw_within_from_Nem: true)*

*Nem_neighborL_min: 1*

*Nem_neighborL_max: 50*

*Nem_neighborR_min: 1*

*Nem_neighborR_max: 50*

*# Bridge migration (Model 2) from Nem (per direction).*

*# If true, bridge m are computed as Nem / Ne_destination; otherwise use m priors above.*

draw_bridge_from_Nem: true

Nem_bridge_L2R_min: 1

Nem_bridge_L2R_max: 25

Nem_bridge_R2L_min: 1

Nem_bridge_R2L_max: 25

### Safety cap on migration probabilities (protects against tiny Ne with large Nem -> m>1)

mig_cap: 0.25

### ---- Effective population size (Ne) priors ----

### Model 1

prior_NeA_min: 1.0e2

prior_NeA_max: 2.0e5

prior_NeD1_min: 1.0e2

prior_NeD1_max: 2.0e4

### Models 2 & 3

prior_NeA2_min: 1.0e2

prior_NeA2_max: 2.0e5

prior_NeLlin_min: 1.0e2     # Left lineage (before splitting into demes)

prior_NeLlin_max: 2.0e5

prior_NeRlin_min: 1.0e2     # Right lineage (before splitting into demes)

prior_NeRlin_max: 2.0e5

prior_NeDL_min: 1.0e2      # Left demes (after T2L)

prior_NeDL_max: 2.0e4

prior_NeDR_min: 1.0e2      # Right demes (after T2R)

prior_NeDR_max: 2.0e4

### ---- How many prior draws per model (rows) ----

### Set to 0 to skip a model entirely.

n_sims1: 10000         # Model 1

n_sims2: 50000           # Model 2 (Secondary Contact)

n_sims3: 10000           # Model 3 (No Secondary Contact)

### ---- Output ----

outdir: All_models_70kLoci  # If null, an auto-named folder is created under CWD

***Supplementary Table S3:*** **$Q_\pi$ estimates**. *Medians and 97% Confidence Intervals (CI) for autosomal, x chromosome nucleotide diversity and their ratios ($\pi_X/\pi_{Autosome}$) for each Carcharhinus amblyrhynchos population and three reference species. Estimates are from 100 site frequency spectra bootstraps, showing values below the expectation of 0.75.*

| Population | Autosome$\pi$ | X$\pi$ | Q$\pi$ | Region |
|---|---|---|---|---|
| *Carcharhinus acronotus* | $9.17x\ 10^{-4}$ [$9.03x10^{-4}$; $9.31x10^{-4}$] | $6.69x10^{-4}$ [$5.08x10^{-4}$; $8.74x10^{-4}$] | 0.73 [0.56; 0.95] | Reference species |
| *Carcharhinus limbatus* | $2.87x\ 10^{-3}$ [$2.87x10^{-3}$; $2.88x10^{-3}$] | $1.27x10^{-3}$ [$1.15x10^{-3}$; $1.38x10^{-3}$] | 0.44 [0.40; 0.48] | Reference species |
| *Galeardo cuvier* | $2.63x\ 10^{-3}$ [$2.59x10^{-3}$; $2.67x10^{-3}$] | $9.8x10^{-4}$ [$6.74x10^{-4}$; $1.29x10^{-3}$] | 0.37 [0.25; 0.49] | Reference species |
| *Juan* | $1.72x\ 10^{-3}$ [$1.71x10^{-3}$; $1.73x10^{-3}$] | $4.13x10^{-4}$ [$3.59x10^{-4}$; $4.76x10^{-4}$] | 0.24 [0.20; 0.28] | Western Indian Ocean |
| *Zelee* | $1.66x\ 10^{-3}$ [$1.65x10^{-3}$; $1.67x10^{-3}$] | $4.57x10^{-4}$ [$3.85x10^{-4}$; $5.5x10^{-4}$] | 0.27 [0.23; 0.33] | Western Indian Ocean |
| *Chagos* | $1.37x\ 10^{-3}$ [$1.35x10^{-3}$; $1.38x10^{-3}$] | $2.45x10^{-4}$ [$1.41x10^{-4}$; $3.95x10^{-4}$] | 0.18 [0.10; 0.29] | Central Indian Ocean |
| *Maldives* | $2.15x\ 10^{-3}$ [$2.13x10^{-3}$; $2.16x10^{-3}$] | $6.4x10^{-4}$ [$5.59x10^{-4}$; $7.14x10^{-4}$] | 0.30 [0.26; 0.33] | Central Indian Ocean |
| *Ningaloo* | $2.57x\ 10^{-3}$ [$2.55x10^{-3}$; $2.59x10^{-3}$] | $1.03x10^{-3}$ [$8.05x10^{-4}$; $1.28x10^{-3}$] | 0.4 [0.31; 0.49] | Eastern Indian Ocean |
| *Rowley* | $2.56x\ 10^{-3}$ [$2.53x10^{-3}$; $2.58x10^{-3}$] | $9.71x10^{-4}$ [$8.22x10^{-4}$; $1.2x10^{-3}$] | 0.38 [0.32; 0.47] | Eastern Indian Ocean |
| *Scott* | $2.57x\ 10^{-3}$ [$2.55x10^{-3}$; $2.6x10^{-3}$] | $1.04x10^{-3}$ [$8.16x10^{-4}$; $1.3x10^{-3}$] | 0.41 [0.32; 0.51] | Eastern Indian Ocean |
| *Bampton* | $3.07x\ 10^{-3}$ [$3.06x10^{-3}$; $3.08x10^{-3}$] | $1.25x10^{-3}$ [$1.16x10^{-3}$; $1.35x10^{-3}$] | 0.40 [0.38; 0.44] | Western Pacific Ocean |
| *Chesterfield* | $2.45x\ 10^{-3}$ [$2.42x10^{-3}$; $2.48x10^{-3}$] | $9.87x10^{-4}$ [$7.20x10^{-4}$; $1.22x10^{-3}$] | 0.4 [0.29; 0.50] | Western Pacific Ocean |
| *Entrecasteaux* | $2.47x\ 10^{-3}$ [$2.44x10^{-3}$; $2.49x10^{-3}$] | $9.62x10^{-4}$ [$8.09x10^{-4}$; $1.21x10^{-3}$] | 0.39 [0.33; 0.49] | Western Pacific Ocean |
| *GLN* | $2.46x\ 10^{-3}$ [$2.43x10^{-3}$; $2.49x10^{-3}$] | $1.07x10^{-3}$ [$8.97x10^{-4}$; $1.36x10^{-3}$] | 0.44 [0.36; 0.55] | Western Pacific Ocean |
| *Grand Astrolabe* | $2.46x\ 10^{-3}$ [$2.44x10^{-3}$; $2.48x10^{-3}$] | $1.03x10^{-3}$ [$7.76x10^{-4}$; $1.27x10^{-3}$] | 0.42 [0.31; 0.51] | Western Pacific Ocean |
| *Indo* | $2.59x\ 10^{-3}$ [$2.57x10^{-3}$; $2.62x10^{-3}$] | $9.64x10^{-4}$ [$7.46x10^{-4}$; $1.14x10^{-3}$] | 0.37 [0.29; 0.44] | Western Pacific Ocean |

| | | | | |
|---|---|---|---|---|
| *Matthew* | $2.34 \times 10^{-3}$ [$2.32 \times 10^{-3}$; $2.36 \times 10^{-3}$] | $1.08 \times 10^{-3}$ [$8.32 \times 10^{-4}$; $1.42 \times 10^{-3}$] | 0.46 [0.35; 0.60] | Western Pacific Ocean |
| *Niku* | $3.07 \times 10^{-3}$ [$3.06 \times 10^{-3}$; $3.08 \times 10^{-3}$] | $1.25 \times 10^{-3}$ [$1.16 \times 10^{-3}$; $1.35 \times 10^{-3}$] | 0.41 [0.38; 0.44] | Western Pacific Ocean |
| *North GBR* | $2.55 \times 10^{-3}$ [$2.53 \times 10^{-3}$; $2.58 \times 10^{-3}$] | $1.09 \times 10^{-3}$ [$8.20 \times 10^{-4}$; $1.34 \times 10^{-3}$] | 0.43 [0.32; 0.52] | Western Pacific Ocean |
| *South GBR* | $2.44 \times 10^{-3}$ [$2.41 \times 10^{-3}$; $2.46 \times 10^{-3}$] | $1.02 \times 10^{-3}$ [$7.84 \times 10^{-4}$; $1.3 \times 10^{-3}$] | 0.4 [0.32; 0.53] | Western Pacific Ocean |
| *South NC* | $2.44 \times 10^{-3}$ [$2.41 \times 10^{-3}$; $2.46 \times 10^{-3}$] | $9.45 \times 10^{-4}$ [$7.03 \times 10^{-4}$; $1.23 \times 10^{-3}$] | 0.39 [0.29; 0.50] | Western Pacific Ocean |
| *Walpole* | $2.39 \times 10^{-3}$ [$2.36 \times 10^{-3}$; $2.41 \times 10^{-3}$] | $1.03 \times 10^{-3}$ [$8.04 \times 10^{-4}$; $1.33 \times 10^{-3}$] | 0.43 [0.34; 0.56] | Western Pacific Ocean |
| *Fakarava* | $3.43 \times 10^{-3}$ [$3.42 \times 10^{-3}$; $3.44 \times 10^{-3}$] | $1.26 \times 10^{-3}$ [$1.16 \times 10^{-3}$; $1.34 \times 10^{-3}$] | 0.37 [0.35; 0.40] | Central Pacific Ocean |
| *Palmyra* | $3.32 \times 10^{-3}$ [$3.32 \times 10^{-3}$; $3.33 \times 10^{-3}$] | $1.35 \times 10^{-3}$ [$1.27 \times 10^{-3}$; $1.41 \times 10^{-3}$] | 0.41 [0.38; 0.42] | Central Pacific Ocean |

***Supplementary Table S4: Fst and $Q_{FST}$ bootstrap estimates****. Medians and 97% Confidence Intervals (CI) are reported for autosomal (A) Fst, X chromosome Fst and their ratio (-log(1-AFst)/-log(1-XFst)) for both datasets comparing the Central Indian Ocean (CIO) and Western Indian Ocean (WIO) to other regions – the Eastern Indian Ocean and the Western Pacific populations. Comparisons were either between (i.e. Indian Ocean vs Pacific) or within respective regions. All $Q_{FST}$ values show strong departure from the neutral expectation of 0.75. Estimates are from 1000 bootstraps with population pair replacement to account for nonindependence of pairwise estimates. Medians in bold indicate significantly different statistic comparisons between regional comparisons.*

| *Statistic* | *Region* | *Regional comparison* | *CI lower* | *median* | *CI upper* |
|---|---|---|---|---|---|
| *$AF_{ST}$* | CIO | between | 0.37 | **0.40** | 0.42 |
| | CIO | within | 0.01 | **0.02** | 0.02 |
| | WIO | between | 0.33 | **0.33** | 0.35 |
| | WIO | within | 0.02 | **0.04** | 0.05 |
| *$Q_{FST}$* | CIO | between | 0.51 | 0.57 | 0.65 |
| | CIO | within | 0.35 | 0.47 | 0.64 |
| | WIO | between | 0.53 | **0.59** | 0.63 |
| | WIO | within | 0.83 | **0.92** | 1.05 |
| *$XF_{ST}$* | CIO | between | 0.56 | **0.59** | 0.60 |
| | CIO | within | 0.03 | **0.04** | 0.06 |
| | WIO | between | 0.48 | **0.50** | 0.53 |
| | WIO | within | 0.02 | **0.04** | 0.05 |

***Supplementary Table S5: Estimates, priors, and Coefficient of determination ($Q^2$) for all parameters*.**

| Parameter | Expectation | Median | Quantile (0.025) | Quantile (0.975) | Prior bounds | $Q^2$ |
|---|---|---|---|---|---|---|
| $T_1$ | 95204 | 77898 | 20956 | 255864 | 100 - 500 000 | 0.674 |
| $T_2L$ | 64846 | 52855 | 17456 | 185760 | 0.1 - 0.9 of $T_1$ | 0.635 |
| $T_2R$ | 61655 | 49144 | 10202 | 149280 | 0.1 - 0.9 of $T_1$ | 0.630 |
| $T_3$ | 24655 | 23582 | 4504 | 42395 | 0.1 - 0.9 of $T_2$ | 0.893 |
| $N_eA$ | 35499 | 28838 | 5208 | 120540 | 100 - 200 000 | 0.624 |
| $N_eL$ | 90391 | 89781 | 2733 | 187102 | 100 - 50 000 | 0.076 |
| $N_eR$ | 102243 | 90835 | 13430 | 193759 | 100 - 50 000 | 0.063 |
| $N_eDL$ | 1611 | 1629 | 996 | 1883 | 100 - 20 000 | 0.905 |
| $N_eDR$ | 5151 | 5047 | 3951 | 8262 | 100 - 20 000 | 0.931 |
| $m_nL$ (%) | 1.75e-02 | 1.58e-02 | 1.17e-03 | 6.10e-02 | $1^{e-4}$ - 0.05 | 0.770 |
| $m_nR$ (%) | 5.87e-03 | 5.88e-03 | 1.99E-03 | 1.03E-02 | $1^{e-4}$ - 0.05 | 0.974 |
| $m_{bLtR}$ (%) | 1.59e-03 | 1.59E-03 | 2.97E-04 | 3.95E-03 | $1^{e-6}$ - 0.02 | 0.742 |
| $m_{bRtL}$ (%) | 1.03E-02 | 1.07E-02 | 1.52E-03 | 1.74E-02 | $1^{e-6}$ - 0.05 | 0.641 |

#### Supplementary Figures

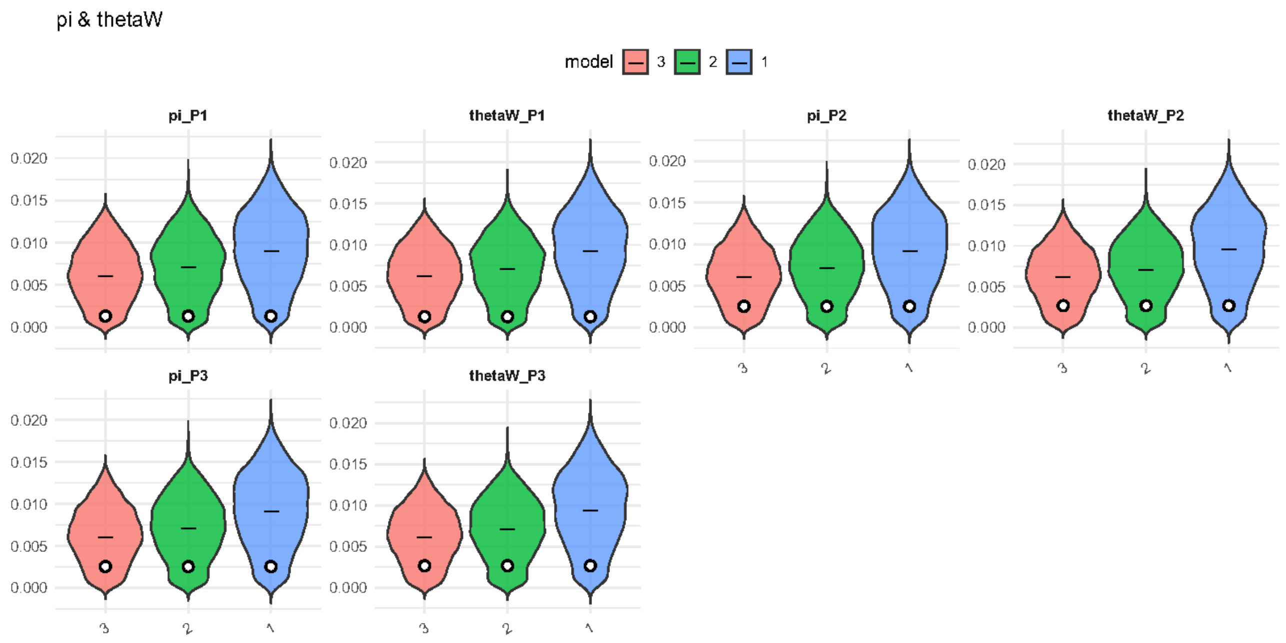


***Supplementary Figure S1: π and theta from ABC simulation****. Violin plots show distribution and mean of stats from the simulations for each model, observed values are noted as dots. Model 1=SST, Model 2 =HIERSC, Model 3 = HIER.*

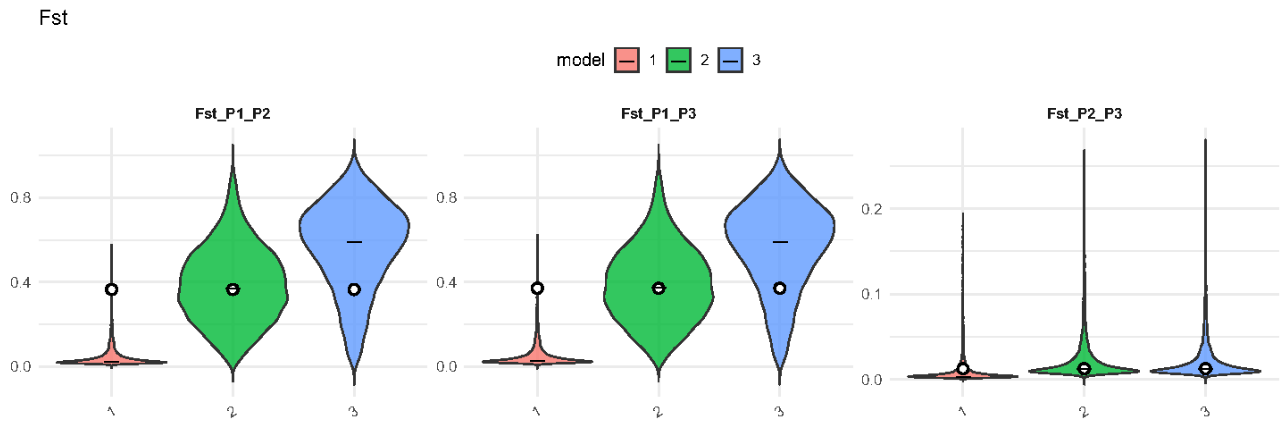


***Supplementary Figure S2****: Pairwise Fst estimates from ABC. Violin plots show distribution and mean of stats from the simulations for each model, observed values are noted as dots. Model 1=SST, Model 2 =HIERSC, Model 3 = HIER.*

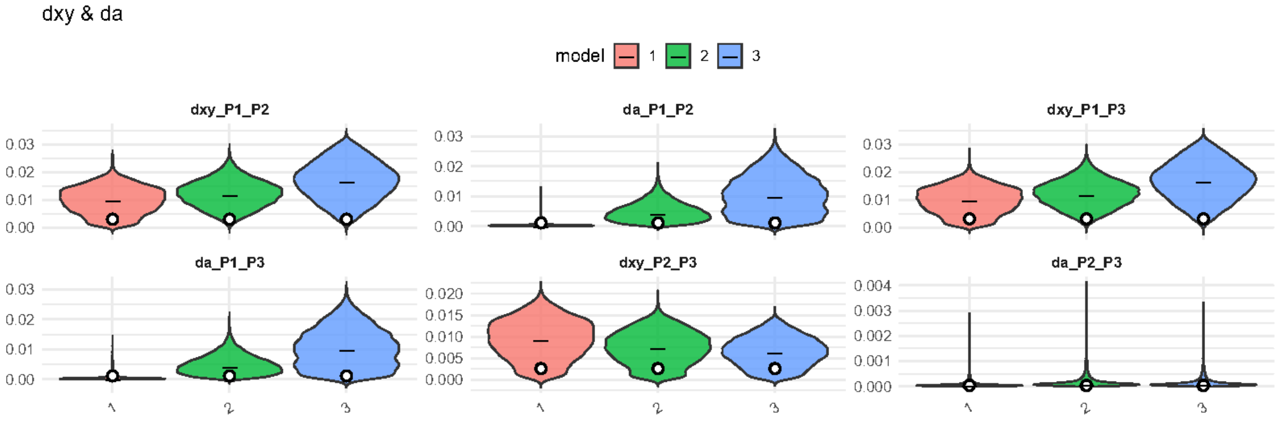


***Supplementary Figure S3:*** *Pairwise dxy and da estimates from ABC. Violin plots show distribution and mean of stats from the simulations for each model, observed values are noted as dots. Model 1=SST, Model 2 =HIERSC, Model 3 = HIER.*

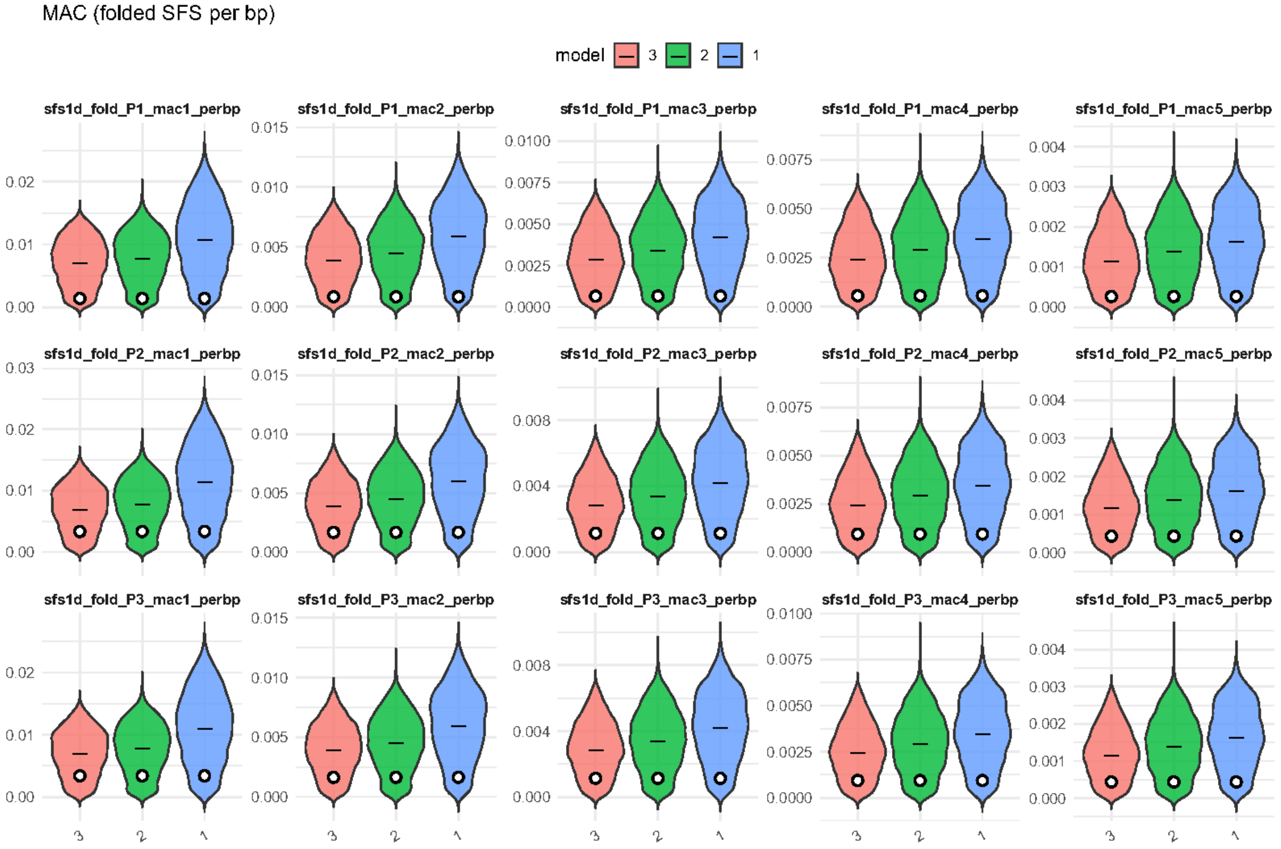


***Supplementary Figure S4:*** *Minor allele counts (MAC). Violin plots show distribution and mean of stats from the simulations for each model, observed values are noted as dots. MAC are for each folded SFS bins for the three populations (n = 5 per population). Model 1=SST, Model 2 =HIERSC, Model 3 = HIER.*

Tajima's D

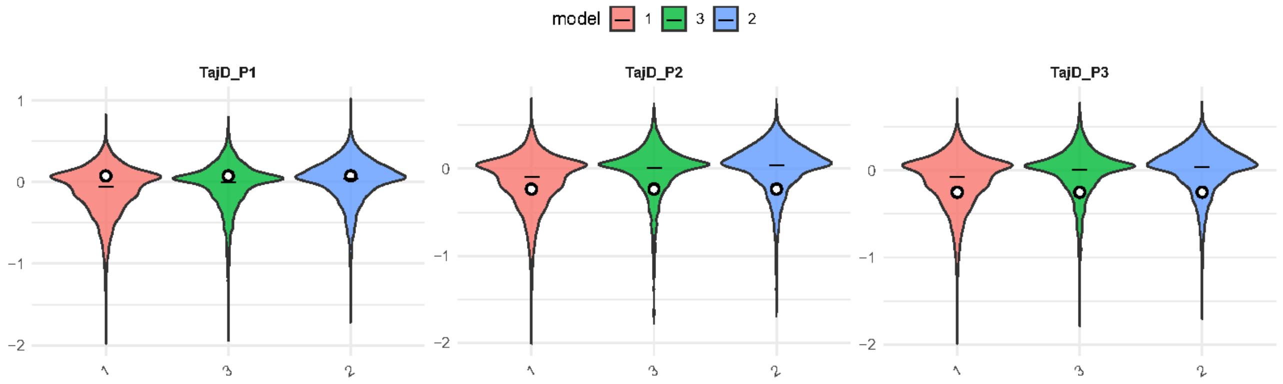


***Supplementary Figure S5**: Tajima's D estimates from ABC. Violin plots show distribution and mean of stats from the simulations for each model, observed values are noted as dots. Model 1=SST, Model 2 =HIERSC, Model 3 = HIER.*

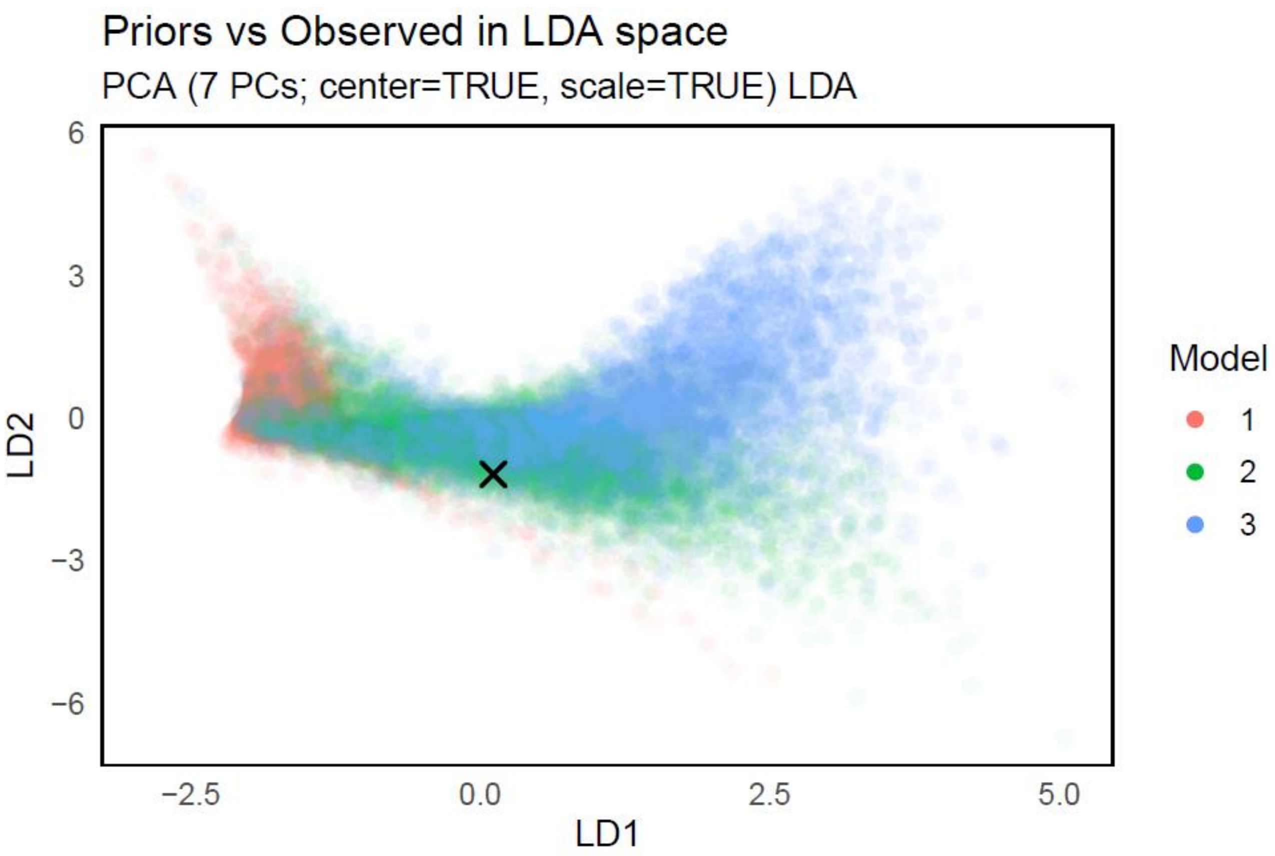


***Supplemtary Figure S6**: Linear Discriminant Analysis of Simulated (colored cicrles) and Observed (X symbol) Summary Statistics for ABC Model Selection. Linear discriminant analysis (LDA) of summary statistics obtained from coalescent simulations under three tested demographic models (1= SST, 2=$HIER_{SC}$, 3=HIER).*

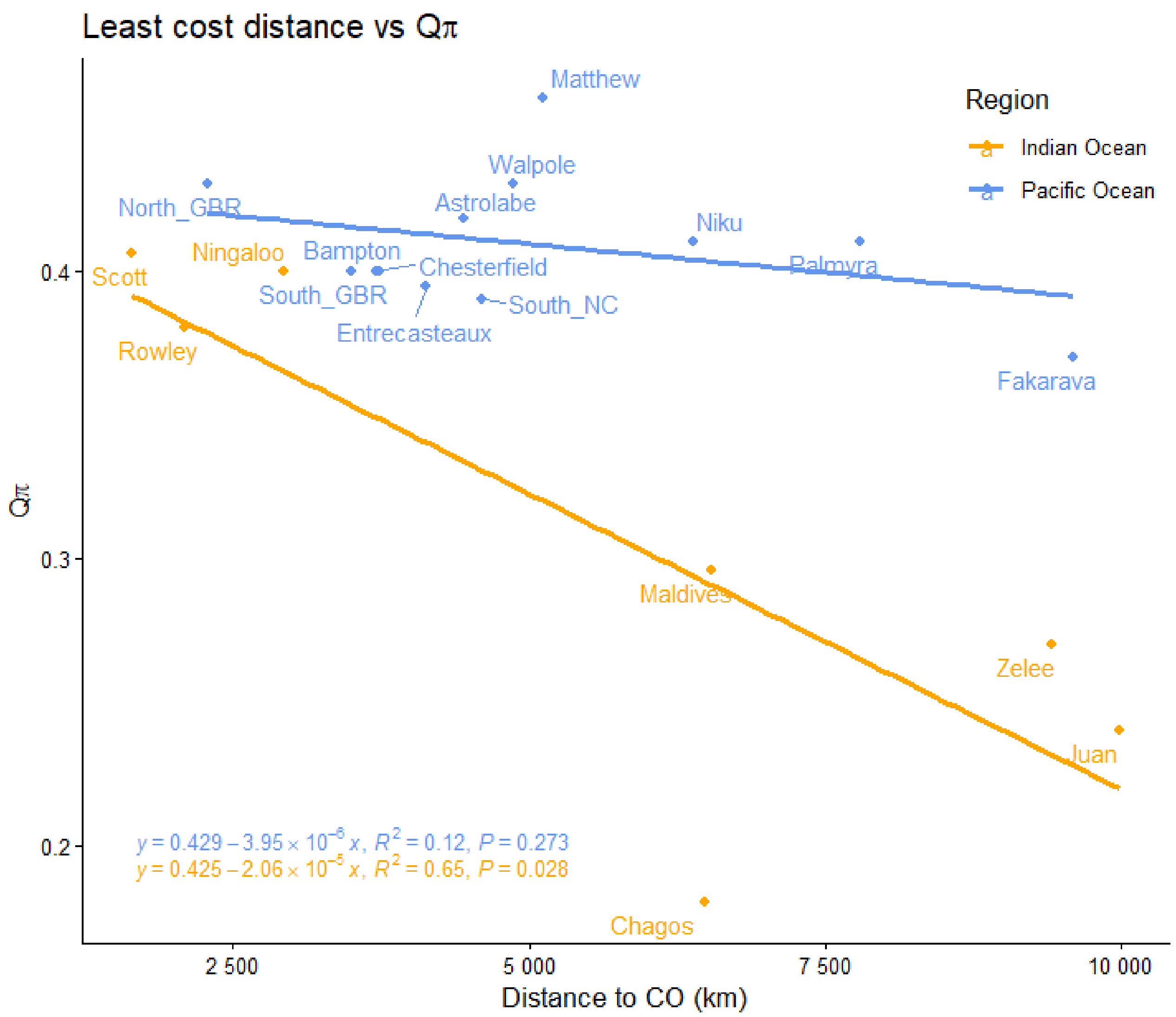


***Supplementary Figure S7: Relationship between $Q_\pi$ and distance to approximate Centre of Origin (CO).*** *For Indian Ocean (orange) and Pacific Ocean (blue) populations, nucleotide diversity ratios ($\pi_X$: $\pi_{autosome}$) are regressed with lest cost distance to approximate centre of past range expansion origin in Indonesia (sample Indo). Distances were calculated in R with marmap (v.1.0.12).*

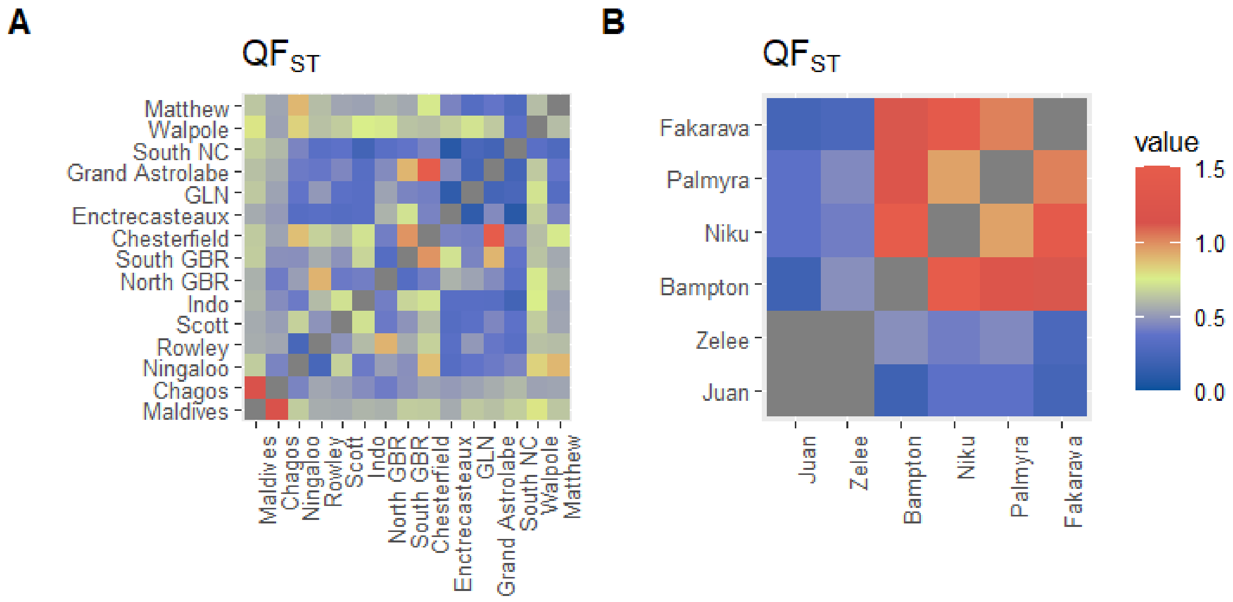


***Supplementary Figure S8: $QF_{ST}$ matrices.*** *Autosome:X Fst ratios for the DArTseq (a) and RADSeq datasets (b). Pseudoautosomal region from the X chromosome and $QF_{ST}$ estimates above 1.5 are removed.*

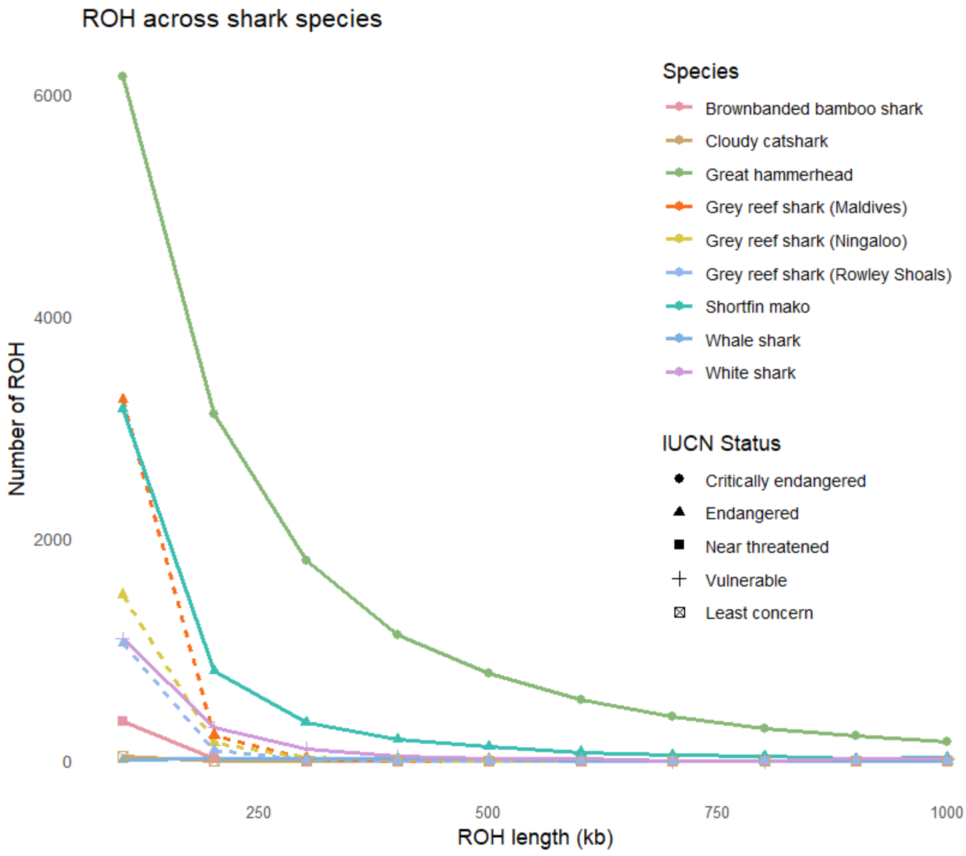


***Supplementary Figure S9: ROH across shark species.*** *Length and number of ROH identified for 24 largest scaffolds of each species as per Stanhope et al., 2023, including estimates from grey reef sharks (green). Shapes represent respective IUCN statuses.*
